# Cardiac nitric oxide synthase 1 worsens heart failure with preserved ejection fraction through S-nitrosylation of histone deacetylase 2

**DOI:** 10.1101/708297

**Authors:** Somy Yoon, Mira Kim, Hangyeol Lee, Gaeun Kang, Kwang-Il Nam, Hyun Kook, Gwang Hyeon Eom

## Abstract

Although the clinical importance of heart failure with preserved ejection fraction (HFpEF), which makes up half of heart failure, has been extensively explored, most therapeutic regimens, including nitric oxide (NO) donors, lack therapeutic benefit^1-12^. Here we report that neuronal nitric oxide synthase (nNOS, also known as NOS1) induces HFpEF by S-nitrosylation of histone deacetylase 2 (HDAC2). HFpEF animal models—SAUNA (SAlty drinking water/Unilateral Nephrectomy/Aldosterone)^13,14^ and mild transverse aortic constriction (TAC) mice^14,15^—showed increased nNOS expression and NO production, which resulted in the S-nitrosylation of HDAC2. HFpEF was alleviated in S-nitrosylation-dead HDAC2 knock-in mice. Pharmacologic intervention by either nNOS inhibition or HDAC2 denitrosylation attenuated HFpEF. Our observations are the first to demonstrate a completely new mechanistic aspect in HFpEF, which may provide a novel therapeutic approach to HFpEF. In addition, our results provide evidence for why conventional NO-enhancement trials have not been effective for improving HFpEF.

## Introduction

Heart failure refers to an insufficient circulation of blood to peripheral organs. The well-known type is systolic heart failure, the characteristics of which are a loss of effective myocardium for contraction^16,17^. In recent decades, in part due to the prevalence of metabolic diseases such as diabetes, the patient population with HFpEF has grown, with a parallel increase in the clinical and social burdens of this disease^5,6,8,10^. To date, 50% of total heart failure is regarded as HFpEF^6^. Unfortunately, none of the therapies that have been proven effective for heart failure with reduced ejection fraction (HFrEF) have been proven beneficial for increasing the survival rate of patients with HFpEF^2,11,12,18-21^. It is surprising that even NO donors, which are potent vasodilators, do not have any benefit in the treatment of HFpEF^2,11^, and this raises fundamental questions as whether our strategy of replenishing NO is appropriate. Indeed, in the current work, we propose that NOS/NO-mediated nitrosylation of protein may worsen HFpEF, suggesting critical reconsideration of the therapeutic effect of NO.

## Results

Although no perfect mouse model for HFpEF is available^14^, we applied the SAUNA model (Methods, Fig. 1a), which is known to be most similar to human HFpEF^13,14^. At 30 days after initiation of the protocol, we measured cardiac function: parameters of exercise capacity such as ejection fraction (EF), early diastole (E), and mitral valve annulus movement (E’); and performance on a rotarod treadmill test (Methods). Systolic function was well conserved in the SAUNA group. However, the E/E’ ratio was aberrantly increased (Fig. 1b, Extended Data Fig. 1a). Furthermore, locomotive activities were impaired in SAUNA mice (Extended Data Fig. 1c). Thus, we confirmed that the SAUNA model successfully induced HFpEF. As an alternative model, mild TAC (Methods) was introduced. Four weeks after mild TAC, we assessed cardiac function (Extended Data Fig. 1b, 1c, 1d). We found that mild TAC could also induce HFpEF in less than 1 month.

**Fig. 1.**
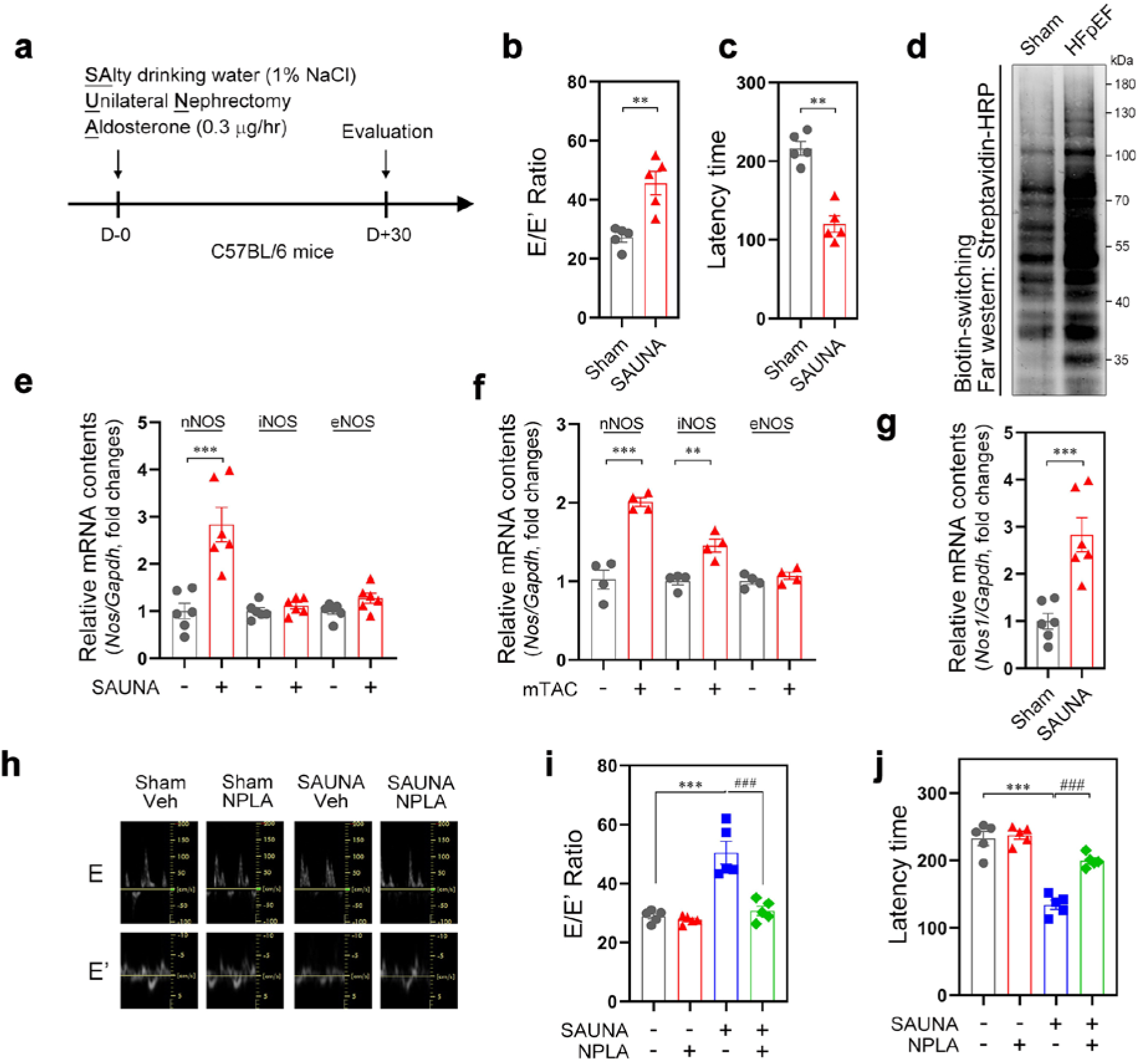
Aberrant activation of neuronal nitric oxide synthase (NOS) in the heart accelerates heart failure with preserved ejection fraction (HFpEF). **a**, Schematic demonstration of the SAUNA (SAlty drinking water/Unilateral Nephrectomy/Aldosterone) model. **b, c**, Changes in SAUNA mice. Elevated mitral early wave (E)/mitral annulus movement (E’) demonstrated diastolic dysfunction (**b**). Decreased latency record in the rotarod treadmill test revealed that SAUNA mice had exercise intolerance (**c**). SAUNA mice are suitable for HFpEF model. **d**, Among the posttranslational modifications of total protein, S-nitrosylation is notably increased in SAUNA heart. 500 μg of heart lysates were labeled by the biotin-switching assay and far western blots with streptavidin-HRP were performed to visualize biotinyl protein. **e-g**, Quantitative real-time PCR to determine transcript amounts of NOS subtype. Transcripts of nNOS (NOS1) were increased in SAUNA-stressed heart (**e**) and both nNOS and iNOS (**f**) were activated in mild TAC. To specify the activation of nNOS in cardiomyocytes, adult cardiomyocyte from SAUNA heart was isolated and transcription activity was determined (**g**). **h, i**, Mitral assay to determine diastolic function. Representative images of Early wave (E) and mitral valve annulus movement (E’). The E/E’ ratio was aberrantly increased in SAUNA mice, which demonstrates moderate to severe diastolic dysfunction. Concomitant treatment with a specific nNOS inhibitor, N(ω)-propyl-L-arginine (NPLA) (50 mg·kg^-1^·day^-1^, every other day), during SAUNA induced normalization both of peak E wave velocity and mitral valve movement. **j**, Exercise capacity was measured by rotarod treadmill test. Latency time on the rolling rod was decreased in SAUNA mice, but was improved by NPLA. **b, c, g,** Unpaired Student’s t-test with 2-tailed mode was used for statistics. *** indicates p<0.001, ** indicates p<0.01. NS, not significant. **i, j**, Multiple comparison was carried out by analysis of variance (ANOVA) followed by Tukey’s post hoc test. ### means p<0.001.

To delineate the mechanism of HFpEF, we measured posttranslational modifications of proteins in both mouse models. Among the posttranslational modifications tested, protein S-nitrosylation was significantly increased (Fig. 1d), whereas no notable alterations in acetylation, methylation, or phosphorylation were observed (Extended Data Fig. 1e, 1f, 1g). NO can be generated by three nitric oxide synthases: nNOS, iNOS2 (NOS2), and eNOS (NOS3)^22^. To understand the source of the NO, we measured transcript amounts of nNOS, iNOS, or eNOS by quantitative real-time PCR. Both nNOS and iNOS were dramatically increased in mild TAC mice, whereas only nNOS was activated in the SAUNA model (Fig. 1e, 1f). We confirmed that nNOS was predominantly increased in cardiomyocytes isolated from SAUNA-treated mice (Methods) (Fig. 1g).

Several pharmacologic agents specifically inhibit nNOS^23-28^. Thus, to confirm the involvement of nNOS, we used N(ω)-propyl-L-arginine (NPLA)^28^ in our HFpEF models. Injection of NPLA every other day (Methods) improved both diastolic dysfunction and exercise capacity (Fig. 1h, 1i, 1j)

Next, we looked for nNOS-specific S-nitrosylation targets in the heart by using a biotin-switching assay^29^ (Methods). Infection of H9c2 cells with adenovirus nNOS-HA generated *de novo* S-nitrosylation (Fig. 2a). By mass spectrometry, we identified that nNOS S-nitrosylated of Hdac2 and Gapdh (Fig. 2a). Incubation of S-nitrosoglutathione (GSNO), a nonselective NO donor, with heart lysates increased S-nitrosylation of both proteins (Extended Data Fig. 2a). S-Nitrosylation of Hdac2 and Gapdh was further confirmed by western blot (Fig. 2b, Extended Data Fig. 2b, 2c).

**Fig. 2.**
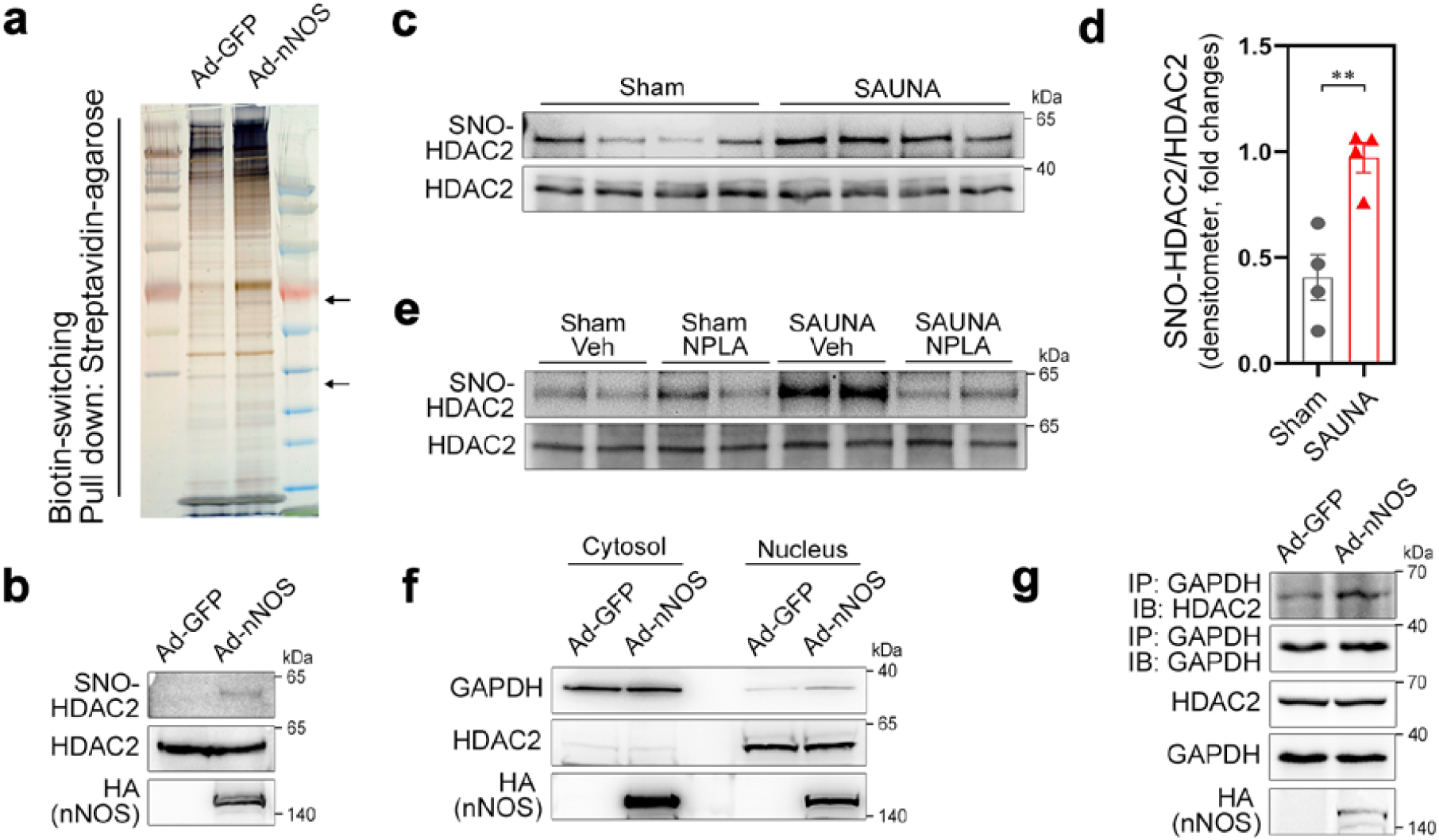
nNOS S-nitrosylates HDAC2 in HFpEF. **a**, Silver staining after biotin-switching assay. H9c2 cells were infected either by adenovirus GFP (Ad-GFP) or adenovirus nNOS (Ad-nNOS), and SNO-cysteine was labeled with biotin. After pulling down with 30 μL of streptavidin-agarose, the biotinyl protein was separated in SDS-PAGE gels and visualized by silver staining. Darkened or *de novo* bands in Ad-nNOS were sequenced by mass spectrometry. Hdac2 (upper arrow) and Gapdh (lower arrow) were identified. **b**, Western blot after biotin-switching assay to further confirm the S-nitrosylation of the identified protein. Infection of Ad-nNOS successfully induced HDAC2 S-nitrosylation in H9c2 cells. **c, d,** S-nitrosylation of HDAC2 in SAUNA heart. Biotin-switching assay to determine S-nitrosylation of HDAC2 in HFpEF. HDAC2 S-nitrosylation was significantly increased in SAUNA heart (**c**). Relative change was measured by densitometer. Unpaired Student’s t-test with 2-tailed mode. ** means p<0.01. (**d**). **e**, Blocking HDAC2 S-nitrosylation by NPLA. Robust S-nitrosylation of HDAC2 by SAUNA stress was completed blocked by NPLA injection. **f, g**, Transnitrosylation of GAPDH to HDAC2. Nitrosative stress derived by nNOS infection induced nuclear targeting of GAPDH after S-nitrosylation (Extended Data Fig. 2b) (**f**). Physical interaction between HDAC2 and GAPDH was increased in the presence of nitric oxide (**g**).

To test whether SAUNA stresses induced S-nitrosylation of Hdac2, we performed the biotin-switching assay with SAUNA heart lysates and observed that S-nitrosylation of Hdac2 was significantly increased (Fig. 2c, 2d). S-Nitrosylation of Hdac2 was also observed in the mild TAC model (Extended Data Fig. 2d). S-Nitrosylation of Hdac2 was nearly completely abolished in SAUNA mice with NPLA administration (Fig. 2e).

nNOS is mainly located in the cytoplasmic membrane, whereas HDAC2 is tethered in the nucleus. Hence, we assumed that nNOS ‘indirectly’ S-nitrosylates Hdac2 by transferring NO from membranous nNOS to nuclear HDAC2. Kornberg et al^30^ reported that GAPDH may work as a shuttling mediator of NO after its S-nitrosylation at Cys150 and after redistribution in the cell. In our experimental models, Gapdh was also S-nitrosylated by nNOS (Fig. 2a), which then resulted in its nuclear redistribution (Fig. 2f). It is also noteworthy that physical interaction between Gapdh and Hdac2 increased in the presence of nNOS (Fig. 2g). When a nonselective nitrosylation-inducer, GSNO, was incubated with heart lysates, Gapdh also physically interacted with Hdac2 (Extended Data Fig. 2e).

Two cysteine residues, Cys262 and Cys274, are known to be responsible for S-nitrosylation of HDAC2^31^. These residues are highly conserved throughout species (Fig. 3a). We generated S-nitrosylation-resistant mutant mice by gene knocking-in technology and substituting those two cysteines with alanines (hereafter, HDAC2 2CA). We could not observe S-nitrosylation of HDAC2 in HDAC2 2CA mouse embryonic fibroblasts even in the presence of adenovirus nNOS-HA (Fig. 3b).

**Fig. 3.**
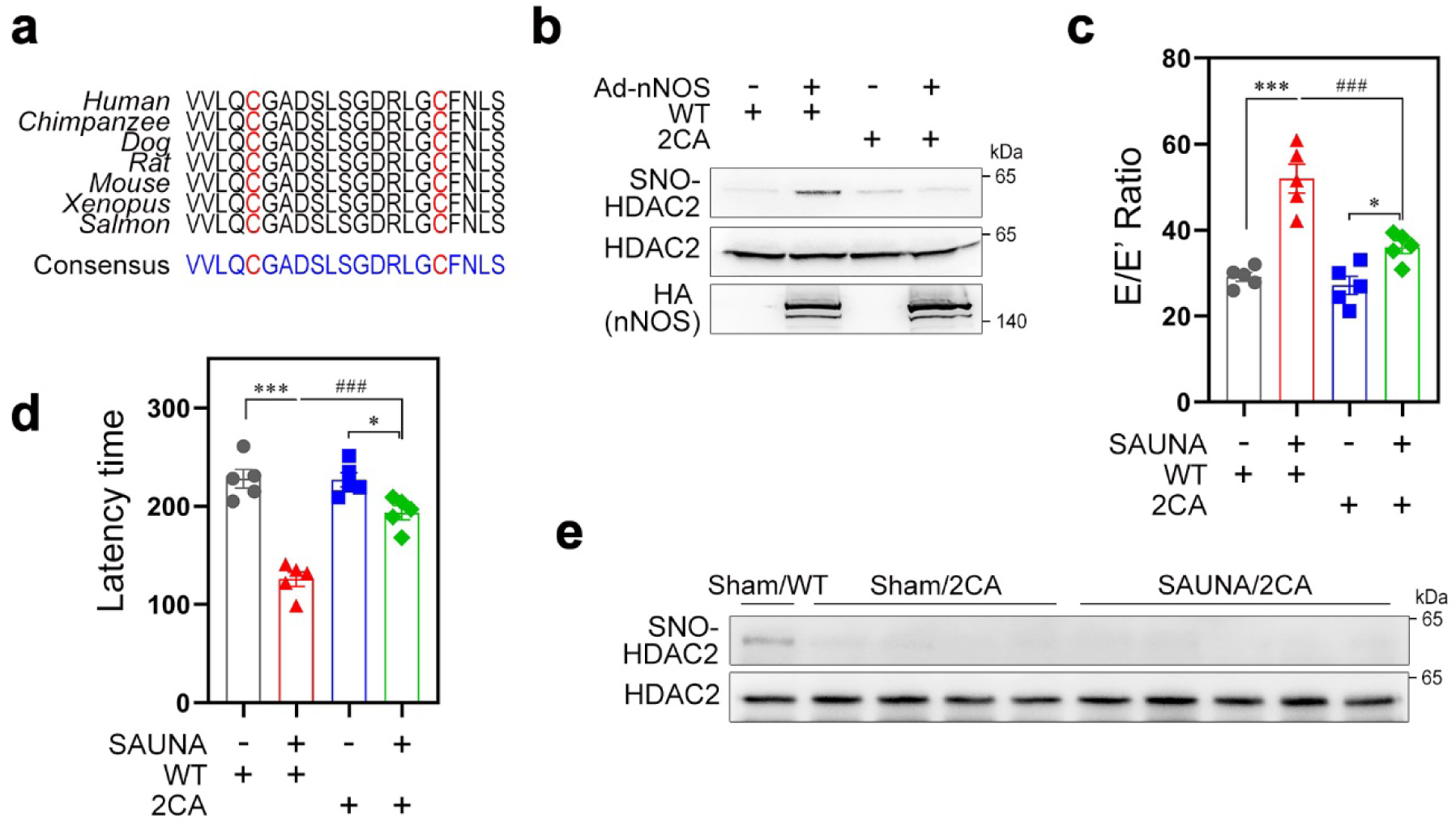
HDAC2 S-nitrosylation undergoes C262 and C274, which exacerbates HFpEF. **a**, Amino acid sequence homology throughout species. Protein sequences flanking C262 and C274 were completely matched in whole species tested. **b**, Biotin-switching assay in S-nitrosylation dead mutant HDAC2 mice (HDAC2 2CA, Methods). Ad-nNOS successfully induced HDAC2 S-nitrosylation, whereas it failed to do in HDAC2 2CA mouse embryonic fibroblast. **c**, Mitral valve assay. SAUNA exacerbates diastolic dysfunction in wild type littermates; on the other hand, SAUNA stresses failed to aggravate diastolic dysfunction in HDAC2 2CA. **d**, Locomotor intolerance. Exercise capacity was reduced by SAUNA in wild type littermates. HDAC2 2CA mice could tolerate SAUNA stress. **c, d**, ANOVA with Tukey’s post hoc. *** and ### indicate p<0.0001. * means p<0.05. **e**, Biotin-switching assay in HDAC2 2CA mice heart. As noted in Fig. **2c**, basal HDAC2 S-nitrosylation was observed in wild type littermates, whereas the S-nitrosylation band was barely detected in HDAC2 2CA mice even under SAUNA stresses. WT indicates wild type littermates and 2CA depicts HDAC2 2CA.

We then applied the SAUNA model to HDAC2 2CA mice. SAUNA failed to exaggerate diastolic dysfunction in HDAC2 2CA mice. The aberrant increase in the E/E’ ratio seen in wild type was not observed in HDAC2 2CA (Fig. 3c). HDAC2 2CA mice also did not develop SAUNA-induced impairment of exercise tolerance (Fig. 3d). As expected, systolic function was well conserved in HDAC2 2CA mice even in SAUNA stresses (Extended Data Fig. 3a). SAUNA-induced HDAC2 S-nitrosylation was not detected in HDAC2 2CA hearts (Fig. 3e). Mild TAC also failed to induce diastolic dysfunction in HDAC2 2CA mice (Extended Data Fig. 3b). Thus, we concluded that HDAC2 2CA mice can tolerate HFpEF.

We next questioned whether S-nitrosylation affects HDAC2 function in HFpEF by measuring HDAC2 activity (Methods). The deacetylase activity of HDAC2 from heart lysates was not changed by GSNO-induced nitrosylation (Extended Data Fig. 3c). Thus, rather than its deacetylase activity, other functions of HDAC2 such as intracellular localization or alterations in complex formation may contribute to the development of HFpEF. This remains to be clarified.

Heretofore, we delineated the roles of nNOS in association with HDAC2 C262/C274 S-nitrosylation in the development of HFpEF. Hence, we questioned whether pharmacologic intervention to reduce the nitrosylation of HDAC2 could also alleviate HFpEF. As reported previously^32^, we observed that NRF2 (also known as NFE2L2) can denitrosylate HDAC2 in cardiac cells. Adenovirus NRF2 significantly activated its target gene, Glutathione S-Transferase Alpha 3 (GSTA3), in H9c2 cells (Extended Data Fig. 4a, 4b). Adenovirus nNOS-HA infection in H9c2 cells successfully induced HDAC2 S-nitrosylation, which was attenuated by infection of adenovirus NRF2 (Fig. 4a).

**Fig. 4.**
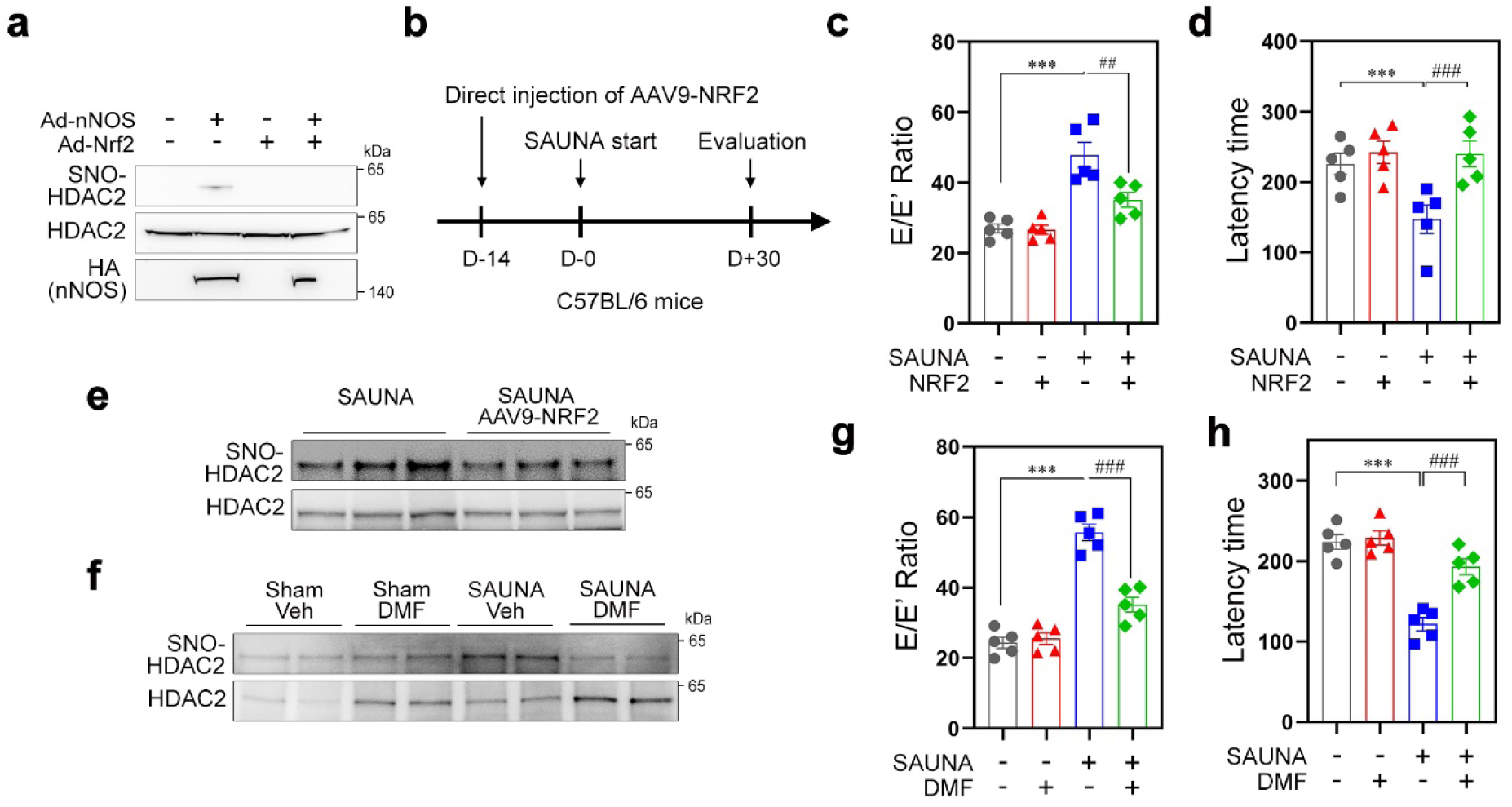
HDAC2 denitrosylation ameliorates HFpEF. **a**, NRF2 denitrosylates HDAC2. Ad-nNOS induced HDAC2 S-nitrosylation, which was completely inhibited by simultaneous infection of adenovirus NRF2. **b**, *in vivo* delivery scheme for overexpression of NRF2 in the adult heart. To allow enough time for activation in the heart, AAV9-NRF2 was directly injected 2 weeks prior to undergoing the SAUNA model (Methods). **c, d**, Overexpression of NRF2 in the heart successfully alleviated diastolic dysfunction (**c**) as well as exercise intolerance (**d**). **e**, AAV9-NRF2 reduced HDAC2 S-nitrosylation in the SAUNA heart. **f, g, h**, dimethyl fumarate (DMF, 0.5 g·l^-1^) inhibited development of HFpEF through denitrosylation of HDAC2. HDAC2 S-nitrosylation was not detected in DMF-treated SAUNA mice (**f**). Elevation of E/E’ was attenuated (**g**) and exercise ability was also improved by DMF (**h**). **c, d, g, h**, ANOVA with Tukey’s post hoc. ### and *** mean p<0.0001, ## indicates p<0.001.

To test the effect of NRF2-induced denitrosylation on HFpEF, we designed an AAV9-NRF2 delivery scheme (Fig. 4b). Direct cardiac injection of AAV9-NRF2 ameliorated diastolic dysfunction in SAUNA mice (Fig. 4c) without any significant changes in systolic function (Extended Data Fig. 4c). Locomotor intolerance was also improved by cardiac overexpression of NRF2 (Fig. 4d). As expected, S-nitrosylation of HDAC2 was notably attenuated in the AAV9-NRF2 injection group (Fig. 4e). AAV9-NRF2 infection was confirmed by transcription amounts of the target gene, GSTA3 (Extended Data Fig. 4d).

Although the specific mechanism remains unclear, dimethyl fumarate (DMF) has been reported to induce NRF2 transcription^33^. We tested whether DMF could alleviate the HFpEF phenotype (Methods). Daily intake of DMF activated NRF2 and subsequently induced GSTA3 (Extended Data Fig. 4e, 4f). Like AAV9-NRF2, HDAC2 S-nitrosylation was not observed in DMF-administered mice (Fig. 4f). DMF also improved both diastolic dysfunction (Fig. 4g) and exercise tolerance (Fig. 4h), which suggests DMF as a novel therapeutic strategy for HFpEF.

To study whether S-nitrosylation of HDAC2 is also involved in HFrEF, we induced severe TAC in HDAC2 2CA mice or wild type littermates (Methods) and measured survival for 15 weeks. Interestingly, HDAC2 2CA mice showed no survival benefit compared with their wild type littermates in this model (Extended Data Fig. 4g). Severe impairment of EF, a surrogate indicator of HFrEF, was also observed as either in the wild type mice or in DMF-administered mice (Extended Data Fig. 4h). Similarly, DMF did not improve the survival rate of wild type mice exposed to severe TAC. These findings suggested that nitrosative stress specifically induces HFpEF rather than general heart failure.

No established regimens are currently available for the patients with HFpEF^3,6,10,17,34^. Whereas NO has beneficial effects like vasodilation and reduction of afterload in HFrEF, NO donors are not at all effective for improving the survival of HFpEF^2,11^. Although preliminary, our new findings that NO-mediated nitrosylation provokes HFpEF may explain why NO donors do not work in the clinical setting. Our study is not the only work to suggest NO as an inducer of HFpEF; indeed, a recent report also implied NOS/NO-mediated nitrosylation of protein in the development of HFpEF^35^. Thus, it is very likely that S-nitrosylation of key proteins may lead the scientific world to revisit NO as a potential therapeutic target of HFpEF. Of note, unlike in HFrEF, however, inhibition of NOS or removal of S-nitrosylation could be considered in HFpEF.

## Methods

### Antibodies

Antibodies used were as follows: mouse monoclonal anti-HDAC2 (Abcam, 1:5,000, 12169), rabbit polyclonal anti-HDAC2 (Invitrogen, 1:1,000, 51-5100), mouse monoclonal GAPDH (Bio-Rad, 1:1,000, VMA00046), rabbit polyclonal anti-GAPDH (Bio-Rad, 1:1,000, VPA00187), and mouse monoclonal anti-HA (Sigma, 1:30,000, H9658).

### Animal model

The use of animal experiments for disease models was approved by the Chonnam National University Medical School Research Institutional Animal Care and Use Committee (CNU IACUC-H-2019-3). For the HFpEF model, 8-week-old male C57BL/6 mice underwent the SAUNA (SAlty drinking water/Unilateral Nephrectomy/Aldosterone) model. The mice were anesthetized with 2,2,2-tribromoethanol (300 mg·kg^-1^, Sigma, T48402) and placed in the right lateral decubitus position. The left kidney was removed and a micro-osmotic pump (Alzet®, Durect Corp, 1004) containing d-aldosterone (Sigma, 0.30 μg·h^-1^, A9477) was implanted under the back skin. Starting 1 day after the operation, the mice were given drinking water with 1% NaCl for 30 days. Exercise capacity and cardiac function were assessed at 30 days. N(ω)-Propyl-L-arginine (NPLA, Cayman, 80587) (50 mg·kg^-1^·day^-1^, every other day) was injected to inhibit nNOS until the end of the study. To induce Nrf2, DMF (Sigma, 0.5 g·l^-1^ in drinking water) was administered together with drinking water for 30 days. AAV9-NRF2 infection (Vector Biolabs, AAV-216638) was carried out by direct cardiac injection. C57BL/6 mice were anesthetized with 2,2,2-tribromoethanol and maintained with an artificial ventilator. The intercostal space between the 4^th^ and 5^th^ rib was cut and widened. 1×10^12^ genome copies of AAV9-NRF2 was injected in the left ventricular free wall. To guarantee successful expression of infected-AAV9-NRF2, the AAV virus was injected 2 weeks before the SAUNA operation. For an alternative HFpEF animal model, mild TAC was carried out to induce isolated diastolic dysfunction without a concomitant change in systolic function. A partial thoracotomy was performed at the second rib and two loose knots were made between the brachiocephalic artery and the left common carotid artery with a 26-gauge needle. For severe TAC, a 27.5-gauge needle was utilized.

### Adult cardiomyocyte isolation

Adult mouse ventricular cardiomyocytes were obtained from C57BL/6 mouse heart. Mice were injected with 50 units of heparin and were euthanized by cervical dislocation. The heart was quickly harvested and cannulated with calcium-free Tyrode buffer (10 mM HEPES pH 7.4, 137 mM NaCl, 5.4 mM KCl, 1 mM MgCl_2_, 10 mM glucose, 5 mM taurine, and 10 mM 2,3-butanedione monoxime) for 3 min with 100% oxygen. The enzyme digestion was carried out by digestion buffer (calcium-free Tyrode buffer supplemented with hyaluronidase [Worthington, 0.1 mg·ml^-1^, LS005477] and collagenase type B [Roche, 0.35 U·ml^-1^, 11088807001]). The left ventricle was collected after 10 minutes’ digestion and cut into small pieces. Further digestion was performed with gentle stirring for 10 minutes. The cells were allowed to stand briefly for large parts to settle and supernatants were filtered by use of a 100-mm pre-cell strainer. The filtered cells were plated on culture dishes for 2 hours to remove cardiac fibroblasts.

### Biotin-switching assay

Protein S-nitrosylation was measured after biotin-switching^29^ with slight modification. All reactions were carried out in dark amber tubes to avoid UV exposure. Protein lysates were prepared with HENS buffer (250 mM HEPES, pH 7.7, 1 mM EDTA, 0.1 mM neocuproine [Sigma, P9599], 1% SDS) and protease inhibitor cocktail. Lysates were prepared in 200 μL of HENS buffer containing 100 μg (cell lysate) or 500 μg (heart lysate) of protein. Free thiol groups were blocked with blocking solution (250 mM HEPES, pH 7.7, 1 mM EDTA, 0.1 mM neocuproine [Sigma, P9599], 1% SDS, and 25 mM S-methylmethane thiosulfonate [Sigma, 208795]) for 1 hour at room temperature. The blocking step was terminated by adding 1 mL of ice-cold acetone. Protein was precipitated by centrifugation at 6,000 RCF for 15 minutes. After removing acetone, the pellet was dissolved in 170 μL of HENS buffer. S-NO was removed by adding 10 μL of 1 M sodium ascorbate for 20 minutes, and subsequent biotin labeling was carried out by adding 20 μL of 4 mM biotin-HPDP (Thermo, 21341) for 40 minutes. The reaction was stopped by ice-cold acetone precipitation. Biotin-labeled protein was dissolved in 400 μL of HENS buffer and neutralized with 800 μL of 1% NP buffer (1% Igepal CA-630, 50 mM Tris pH 8.0, 150 mM NaCl, 1 mM EDTA) and protease inhibitor cocktail. The biotin-switching protein was collected with 30 μL of streptavidin-agarose beads and continuous rotation at 4°C overnight. The beads were washed three times with 1% NP buffer and the samples were analyzed by SDS-PAGE.

### Echocardiogram

Cardiac function was measured by ultrasonography (General Electric Company, Vivid S5). Mice were anesthetized with 2,2,2-tribromoethanol and checked for lack of response with a light touch. At the papillary muscle level, 2-dimension M-mode was acquired from the parasternal long-axis view or parasternal short-axis view. Ejection fraction was determined by the Teichholz formula: EF(%)=(Vd-Vs)/Vd, where Vd indicates LV volume at end diastole and Vs is at end systole, and Vd=[7/(2.4+LVIDd)]×LVIDd^3, Vs=[7/(2.4+LVIDs)]×LVIDs^3, where LVIDd is LV interventricular dimension at end diastole and LVIDs is at end systole. To assess diastolic function, E, and E’ were measured. After measurement of the parasternal short-axis view, the sonographic probe was tilted 45 degrees to visualize the parasternal 4-chamber view. Mitral E waves were recorded with pulse-wave mode at the mitral valve opening. Mitral annulus movement, also known as E’ wave, was assessed from the medial mitral valve annulus with tissue velocity image mode.

### Genetically engineered mice

HDAC2 S-nitrosylation-dead knock-in mice (HDAC2 2CA) were generated by a commercial company (Toolgen). Mouse HDAC2 genomic sequences flanking cysteine 262 and cysteine 274 were exchanged as follows: TGT GGC GCA GAC TCC CTG TCT GGG GAC AGG CTT GGT TGT to GCT GGA GCC GAT AGC CTT AGC GGA GAT CGC CTGGGA GCT. Protein sequences were not changed except for two targeted cysteines (CGADSLSGDRLGC to AGADSLSGDRLGA). Genotype was determined by T7E1 (NEB, M0302). Direct Sanger sequencing of PCR products was requested to further confirm the genotype of the homozygote. The oligomer set for genotyping was as follows: sense: 5’-TGCTGTCAATTTTCCCATGA-3’, antisense: 5’-AGAGTTTGGCATCGAGTTGG-3’.

### Histone deacetylase activity assay

A commercial kit (HDAC-Glo^™^ 2 Assays, Promega, G9590) was used to measure HDAC2 activity. Heart lysates were prepared with 1% NP buffer without EDTA to avoid zinc chelation. Fifty micrograms of heart lysates were mixed with HDAC2 assay substrate and incubated for 15 minutes at room temperature. To check the S-nitrosylation effect of HDAC2, 500 μM of GSNO (Cayman, 82240) or 500 μM of GSH (L-Glutathione, Cayman, 10007461) was added to the assay buffer. The deacetylase activity of HDAC2 was measured with a luminometer. The vehicle-treated condition was regarded as 1 and fold changes were calculated.

### Quantitative real-time polymerase chain reaction

Total mRNA was extracted with TRIzol (Invitrogen, 15596026). cDNA was synthesized by use of random hexamer (M-MLV reverse transcriptase, Invitrogen, 28025013). Quantitative real-time PCR was carried out by using QuantiTect SYBR Green kits (Qiagen, 204143) with a Rotor-Gene Q (Qiagen). PCR analysis was performed in triplicate and the average was regarded as a single result. The relative contents of mRNA transcripts were normalized to those of Gapdh. Specific oligomer sets were as follows:

Mouse GAPDH, sense: 5’-GCATGGCCTTCCGTGTTCCT-3’, antisense: 5’-CCCTGTTGCTGTAGCCGTAT-3’

Mouse nNOS, sense: 5’-ACTGACACCCTGCACCTGAAGA-3’, antisense: 5’-GTGCGGACATCTTCTGACTTCC-3’

Mouse iNOS, sense: 5’-CAGCTGGGCTGTACAAACCTT-3’, antisense: 5’-CATTGGAAGTGAAGCGGTTCG-3’

Mouse eNOS, sense: 5’-CCTCGAGTAAAGAACTGGGAAGTG-3’, antisense: 5’-AACTTCCTTGGAAACACCAGGG-3’

Mouse GSTA3, sense: 5’-TACTTTGATGGCAGGGGAAG-3’, antisense: 5’-GCACTTGCTGGAACATCAGA-3’

Mouse 18s, sense: 5’-GTAACCCGTTGAACCCCATT-3’, antisense: 5’-CCATCCAATCGGTAGTAGCG-3’

Human NRF2, sense: 5’-CACATCCAGTCAGAAACCAGTGG-3’, antisense: 5’-GGAATGTCTGCGCCAAAAGCTG-3’

Rat NRF2, sense: 5’-AGAAGCACACTGAAGGCACGG-3’, antisense: 5’-GAATGTGTTGGCTGTGCTTTAGGTC-3’

Rat GSTA3, sense: 5’-AGTCCTTCACTACTTCGATGGCAG-3’, antisense: 5’-CACTTGCTGGAACATCAAACTCC-3’

Rat GAPDH, sense: 5’-ATGACATCAAGAAGGTGGTG-3’, antisense: 5’-CATACCAGGAAATGAGCTTG-3’

### Rotarod treadmill test

The locomotor tolerance of mice was measured by using a rotarod system (ENV-577M, Med Associate Inc). Mice were pre-adapted to the rod at a fixed speed of 10 rpm for 5 minutes before measurement. Mice were placed on the round rod (35-mm diameter) with a continuous acceleration from 4 to 40 rpm by 1 rpm per 5 seconds. Motor ability was determined as the time of falling off or passive rolling with grasp rod, up to 5 minutes. The best latency time of 3 tests was used for the study. Time interval between each trial was 30 minutes.

### Statistics

Statistical significance was analyzed with PASW Statistics 25 (SPSS, IBM corp). For two independent groups, two-tailed unpaired Student’s t-test was applied after checking for a normal distribution. In the case of more than two groups, one-way analysis of variance was used. Bonferroni’s post hoc test was applied for multiple comparisons when equal variance was assumed by use of Levene statistics, whereas the Dunnett T3 test was used in case of unequal variance. Survival rate was visualized and calculated by use of Prism 8.0 (GraphPad). Significance was determined at values of p < 0.05.

## Acknowledgments

This work was supported by National Research Foundation of Korea (NRF) grants funded by the Korean government (MSIT) (NRF-2019004521, 20191058992, 2018R1A2B3001503), Republic of Korea.

## Author contributions

S.Y. performed most of the experiments. M.K. performed fractional western blot, immunoprecipitation assays, and quantitative real-time PCR. H.L. carried out the rotarod treadmill test. G.K. statistically analyzed the numeric data. K-I.N. performed histology staining for cardiac geometry. S.Y, H.K., and G.H.E. designed the whole study and wrote the manuscript. H.K. and G.H.E. financially supported the study.

## Extended Data Figure and Figure legends

**Extended Data Fig. 1.**
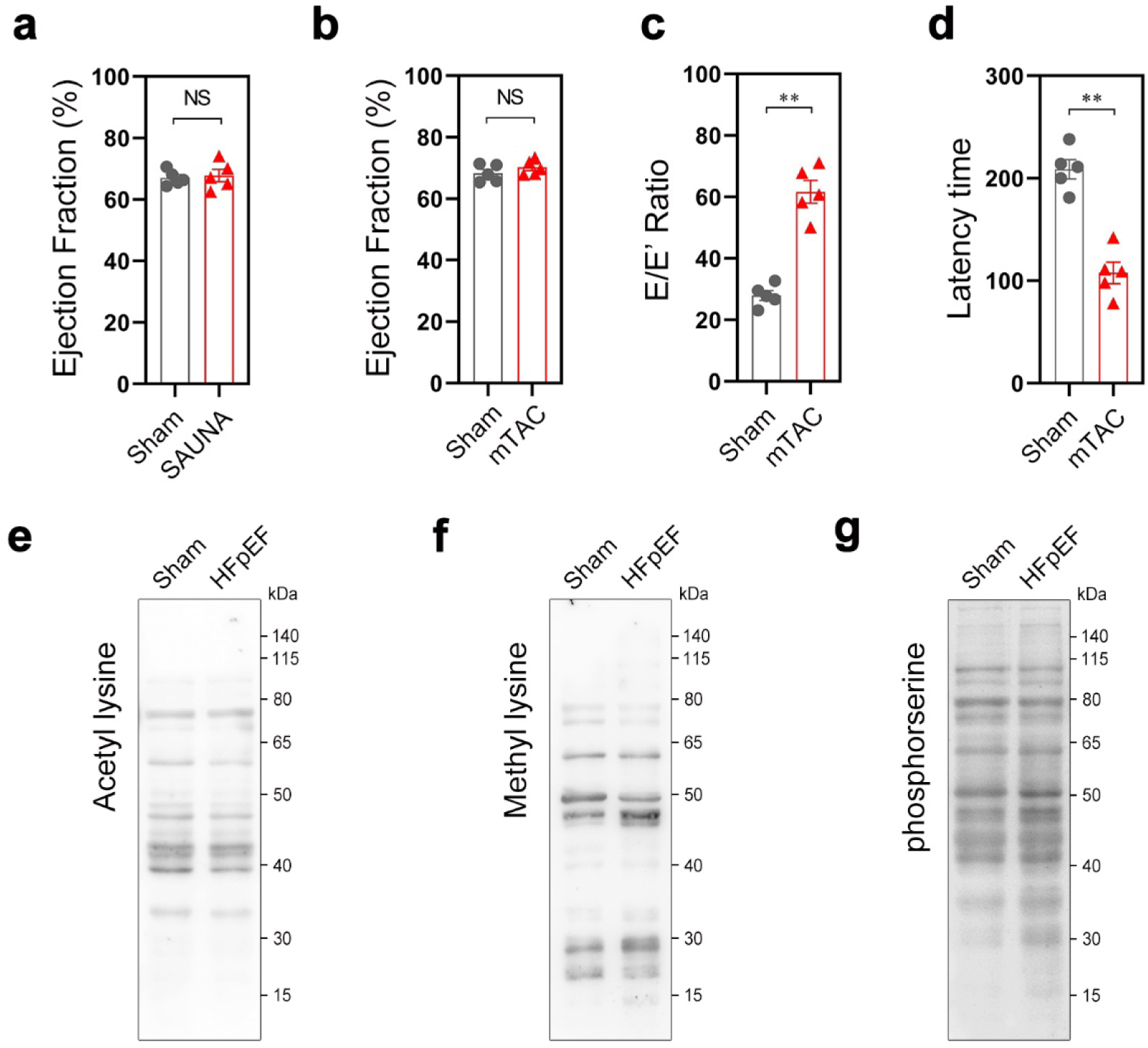
General alteration of HFpEF mice. **a**, Ejection fraction was not changed in SAUNA mice. **b, c, d**, The parameters in mild TAC (mTAC, Methods). Systolic function was not changed in mTAC (**b**). mTAC induced a more severe grade of diastolic dysfunction than did SAUNA (**c**). Exercise capacity was also dramatically reduced in mTAC (d). **e, f, g**, Posttranslational modifications. Total acetylation (**e**), methylation (**f**), and serine phosphorylation (**g**) were not altered in SAUNA heart. **a, b, c, d,** Unpaired Student’s t-test with 2-tailed mode was used for statistics. ** indicates p<0.01. NS, not significant.

**Extended Data Fig. 2.**
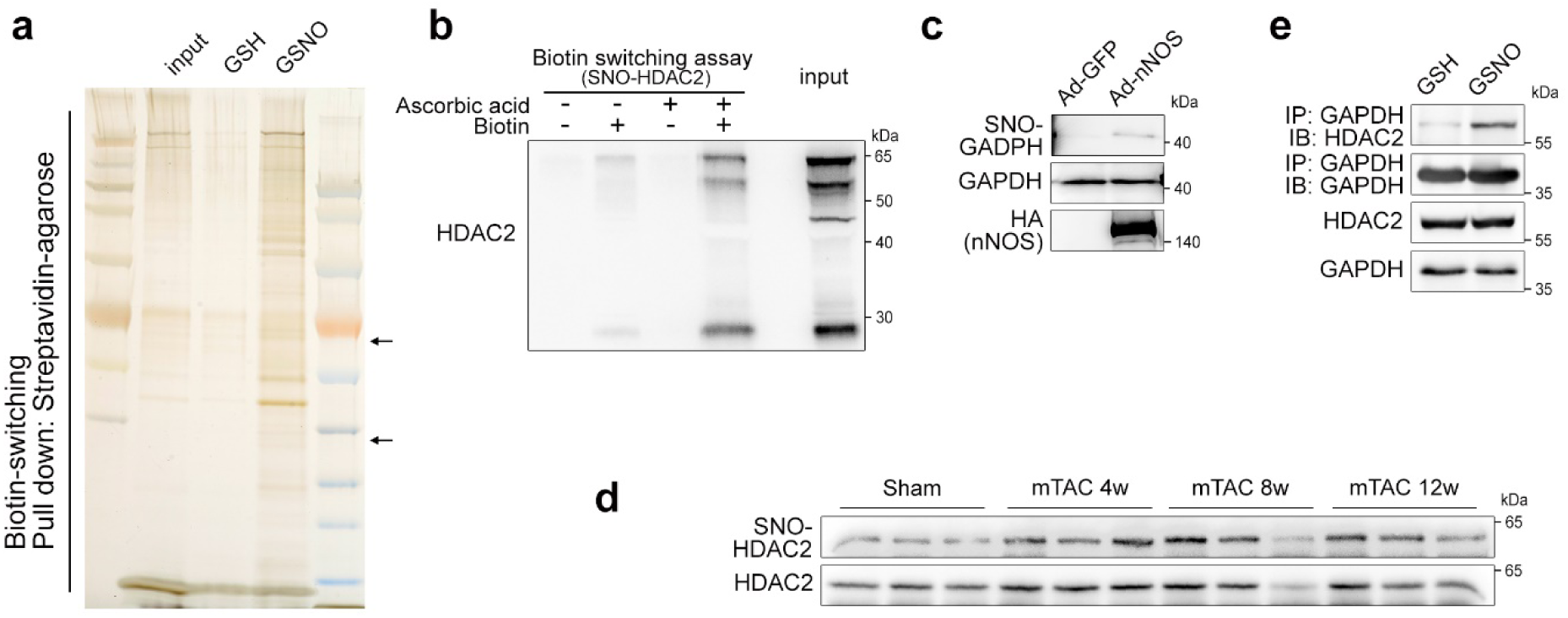
Protein S-nitrosylation in nitrosative stresses. **a**, Silver staining after biotin-switching assay. Note that 500 μM GSNO (S-nitrosoglutathione) generated numerous de novo S-nitrosylation, whereas 500 μM GSH (L-glutathione) decreased it. Hdac2 (upper arrow) and Gapdh (lower arrow) are identified (**a**). **b**, Biotin-switching assay. HDAC2 S-nitroylation was specifically detected when both sodium ascorbate and biotin were added in the reaction. **c**, Infection of adenovirus nNOS induces GAPDH S-nitrosylation. **d**, mTAC S-nitrosylated HDAC2. HDAC2 S-nitrosylation was well detected in 12 weeks after mTAC operation. **e**, In the presence of nitric oxide, physical interaction between HDAC2 and GAPDH was increased.

**Extended Data Fig. 3.**
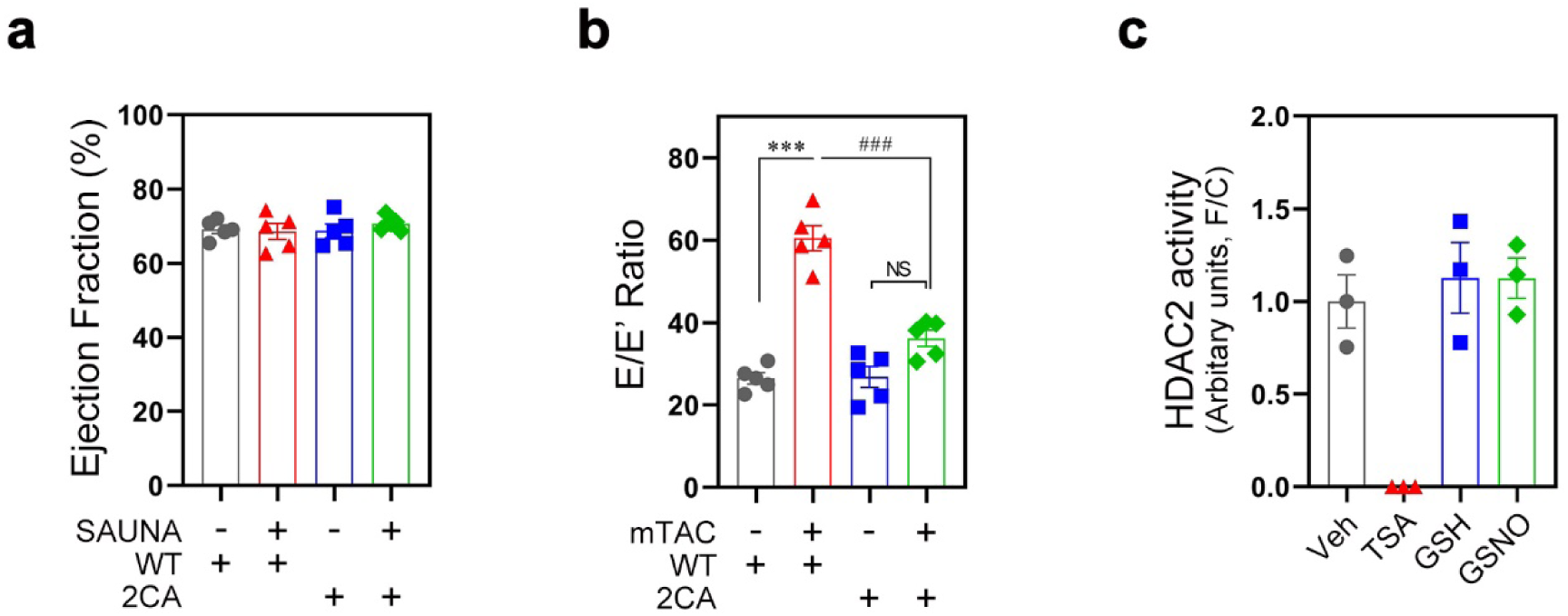
HDAC2 S-nitrosylation at C262 and C274. **a**, Systolic function was well conserved. **b**, Like SAUNA, HDAC2 2CA mice could tolerate mTAC-induced diastolic dysfunction. The E/E’ ratio was not increased in HDAC2 2CA even by mTAC. **c**, HDAC2 activity in the presence of nitric oxide. Antibody-free HDAC2 activity was assessed to investigate the effect of S-nitrosylation on enzyme activity. Equal amounts of heart lysates were mixed with the chemicals noted and deacetylase activity was measured. Trichostatin A (TSA), a potent HDAC inhibitor, was used as the internal control. Note that both GSH and GSNO failed to alter the intrinsic activity of HDAC2 in heart lysates. **a, b, c,** One-way analysis of variance (ANOVA) and Tukey’s post hoc was applied for multiple comparison. *** and ### indicate p<0.0001. NS, not significant.

**Extended Data Fig. 4.**
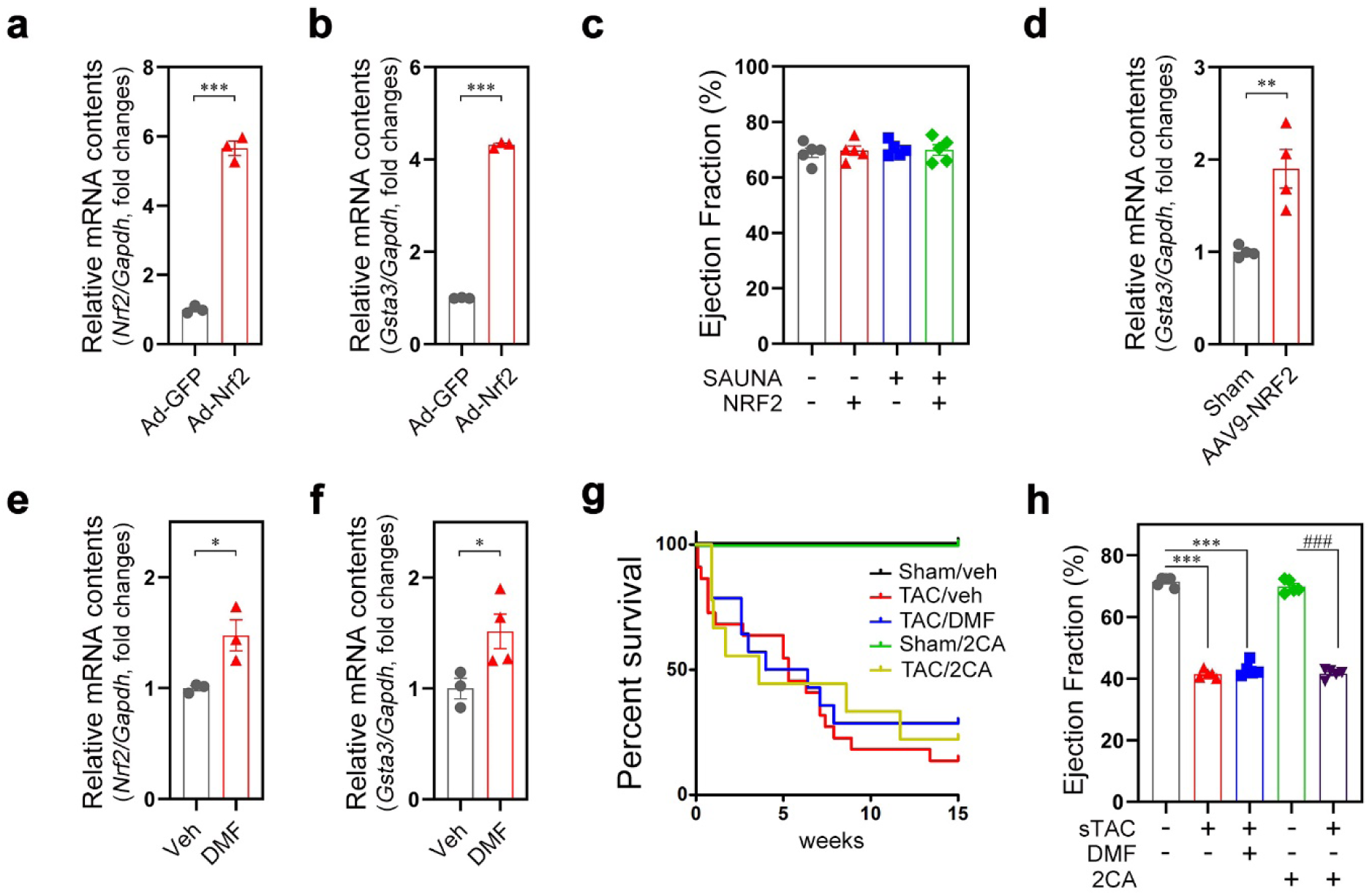
NRF2, a denitrosylase of HDAC2. **a,b**, Adenovirus NRF2 successfully activated transcription both of NRF2 (**a**) and its target gene, GSTA3 (**b**). **c**, Systolic function was not changed in any group. **d,** AAV9-NRF2 infection activated GSTA3 in the heart. **e, f**, Drinking water containing dimethyl fumarate (DMF, 0.5g·l^-1^) activated transcription of NRF2 (**e**) and subsequently its target gene GSTA3 (**f**) in the heart.. **g, h**, Nitric oxide specifically regulates HFpEF. Kaplan-Meier survival curve of systolic heart failure either in HDAC2 2CA or in DMF-treated mice for 15 weeks. HDAC2 2CA revealed that severe TAC (sTAC, Methods) induced non-ischemic systolic heart failure. Similarly, DMF failed to ameliorate survival rate of sTAC. Thereby, the whole life span was not significantly different from that of wild type littermates (**g**) (Mantel-Cox log rank test). Ejection fraction proved systolic heart failure in HDAC2 2CA (**h**). HDAC2 2CA or DMF failed to overcome sTAC-induced HFrEF, which implied that nitrosative stresses are specifically involved in HFpEF rather than general heart failure. **a, b, d, e, f,** Unpaired Student’s t-test with 2-tailed mode. *** indicates p<0.001, ** depicts p<0.01, and * means p<0.05. **c, h,** One-way analysis of variance (ANOVA) and Tukey’s post hoc was applied for multiple comparison. ### indicate p<0.0001.

## References

1. Beldhuis, I.E., et al. Efficacy and Safety of Spironolactone in Patients With HFpEF and Chronic Kidney Disease. JACC Heart Fail 7, 25–32 (2019).

2. Borlaug, B.A., et al. Effect of Inorganic Nitrite vs Placebo on Exercise Capacity Among Patients With Heart Failure With Preserved Ejection Fraction: The INDIE-HFpEF Randomized Clinical Trial. JAMA 320, 1764–1773 (2018).

3. Luscher, T.F. Heart failure subgroups: HFrEF, HFmrEF, and HFpEF with or without mitral regurgitation. Eur Heart J 39, 1–4 (2018).

4. Tanaka, K., et al. Effects of adiponectin on calcium-handling proteins in heart failure with preserved ejection fraction. Circ Heart Fail 7, 976–985 (2014).

5. Borlaug, B.A. & Redfield, M.M. Diastolic and systolic heart failure are distinct phenotypes within the heart failure spectrum. Circulation 123, 2006–2013; discussion 2014 (2011).

6. Dunlay, S.M., Roger, V.L. & Redfield, M.M. Epidemiology of heart failure with preserved ejection fraction. Nat Rev Cardiol 14, 591–602 (2017).

7. Dunlay, S.M., Roger, V.L., Weston, S.A., Jiang, R. & Redfield, M.M. Longitudinal changes in ejection fraction in heart failure patients with preserved and reduced ejection fraction. Circ Heart Fail 5, 720–726 (2012).

8. Mohammed, S.F., et al. Comorbidity and ventricular and vascular structure and function in heart failure with preserved ejection fraction: a community-based study. Circ Heart Fail 5, 710–719 (2012).

9. Mohammed, S.F., et al. Mineralocorticoid accelerates transition to heart failure with preserved ejection fraction via “nongenomic effects”. Circulation 122, 370–378 (2010).

10. Redfield, M.M. Heart Failure with Preserved Ejection Fraction. N Engl J Med 375, 1868–1877 (2016).

11. Redfield, M.M., et al. Isosorbide Mononitrate in Heart Failure with Preserved Ejection Fraction. N Engl J Med 373, 2314–2324 (2015).

12. Redfield, M.M., et al. Effect of phosphodiesterase-5 inhibition on exercise capacity and clinical status in heart failure with preserved ejection fraction: a randomized clinical trial. JAMA 309, 1268–1277 (2013).

13. Hulsmans, M., et al. Cardiac macrophages promote diastolic dysfunction. The Journal of experimental medicine 215, 423–440 (2018).

14. Valero-Munoz, M., Backman, W. & Sam, F. Murine Models of Heart Failure with Preserved Ejection Fraction: a “Fishing Expedition”. JACC Basic Transl Sci 2, 770–789 (2017).

15. Mohammed, S.F., et al. Variable phenotype in murine transverse aortic constriction. Cardiovasc Pathol 21, 188–198 (2012).

16. January, C.T., et al. 2014 AHA/ACC/HRS guideline for the management of patients with atrial fibrillation: a report of the American College of Cardiology/American Heart Association Task Force on practice guidelines and the Heart Rhythm Society. Circulation 130, e199–267 (2014).

17. Ponikowski, P., et al. 2016 ESC Guidelines for the diagnosis and treatment of acute and chronic heart failure: The Task Force for the diagnosis and treatment of acute and chronic heart failure of the European Society of Cardiology (ESC). Developed with the special contribution of the Heart Failure Association (HFA) of the ESC. Eur J Heart Fail 18, 891–975 (2016).

18. Cleland, J.G.F., et al. Beta-blockers for heart failure with reduced, mid-range, and preserved ejection fraction: an individual patient-level analysis of double-blind randomized trials. Eur Heart J 39, 26–35 (2018).

19. Pitt, B., et al. Spironolactone for heart failure with preserved ejection fraction. N Engl J Med 370, 1383–1392 (2014).

20. Shearer, F., Lang, C.C. & Struthers, A.D. Renin-angiotensin-aldosterone system inhibitors in heart failure. Clin Pharmacol Ther 94, 459–467 (2013).

21. Massie, B.M., et al. Irbesartan in patients with heart failure and preserved ejection fraction. N Engl J Med 359, 2456–2467 (2008).

22. Forstermann, U. & Sessa, W.C. Nitric oxide synthases: regulation and function. Eur Heart J 33, 829–837, 837a-837d (2012).

23. Rickard, N.S., Gibbs, M.E. & Ng, K.T. Inhibition of the endothelial isoform of nitric oxide synthase impairs long-term memory formation in the chick. Learn Mem 6, 458–466 (1999).

24. Hah, J.M., Roman, L.J., Martasek, P. & Silverman, R.B. Reduced amide bond peptidomimetics. (4S)-N-(4-amino-5-[aminoakyl]aminopentyl)-N’-nitroguanidines, potent and highly selective inhibitors of neuronal nitric oxide synthase. J Med Chem 44, 2667–2670 (2001).

25. Hicks, C.A., Ward, M.A., Swettenham, J.B. & O’Neill, M.J. Synergistic neuroprotective effects by combining an NMDA or AMPA receptor antagonist with nitric oxide synthase inhibitors in global cerebral ischaemia. Eur J Pharmacol 381, 113–119 (1999).

26. Bland-Ward, P.A. & Moore, P.K. 7-Nitro indazole derivatives are potent inhibitors of brain, endothelium and inducible isoforms of nitric oxide synthase. Life Sci 57, PL131–135 (1995).

27. Rees, D.D., Palmer, R.M., Schulz, R., Hodson, H.F. & Moncada, S. Characterization of three inhibitors of endothelial nitric oxide synthase in vitro and in vivo. Br J Pharmacol 101, 746–752 (1990).

28. Zhang, H.Q., Fast, W., Marletta, M.A., Martasek, P. & Silverman, R.B. Potent and selective inhibition of neuronal nitric oxide synthase by N omega-propyl-L-arginine. J Med Chem 40, 3869–3870 (1997).

29. Zhan, N., et al. S-Nitrosylation Targets GSNO Reductase for Selective Autophagy during Hypoxia Responses in Plants. Mol Cell 71, 142–154 e146 (2018).

30. Kornberg, M.D., et al. GAPDH mediates nitrosylation of nuclear proteins. Nat Cell Biol 12, 1094–1100 (2010).

31. Nott, A., Watson, P.M., Robinson, J.D., Crepaldi, L. & Riccio, A. S-Nitrosylation of histone deacetylase 2 induces chromatin remodelling in neurons. Nature 455, 411–415 (2008).

32. Malhotra, D., et al. Denitrosylation of HDAC2 by targeting Nrf2 restores glucocorticosteroid sensitivity in macrophages from COPD patients. The Journal of clinical investigation 121, 4289–4302 (2011).

33. Hayashi, G., et al. Dimethyl fumarate mediates Nrf2-dependent mitochondrial biogenesis in mice and humans. Hum Mol Genet 26, 2864–2873 (2017).

34. Yancy, C.W., et al. 2017 ACC/AHA/HFSA Focused Update of the 2013 ACCF/AHA Guideline for the Management of Heart Failure: A Report of the American College of Cardiology/American Heart Association Task Force on Clinical Practice Guidelines and the Heart Failure Society of America. Circulation 136, e137–e161 (2017).

35. Schiattarella, G.G., et al. Nitrosative stress drives heart failure with preserved ejection fraction. Nature (2019).

